# Function of basal ganglia in tauG-guiding action

**DOI:** 10.1101/143065

**Authors:** David N. Lee, Apostolos P. Georgopoulos, Gert-Jan Pepping

**Affiliations:** School of Philosophy, Psychology and Language Sciences and Moray House School of Education, University of Edinburgh, St Leonard’s Land, Holyrood Road, Edinburgh EH8 8AQ, UK; Brain Sciences Center, Veterans Affairs Medical Center, One Veterans Drive, Minneapolis, MN 55417, USA; Cognitive Sciences Center, University of Minnesota, Minneapolis, MN 55455, USA; Department of Neuroscience, University of Minnesota Medical School, Minneapolis, MN 55455, USA; Department of Neurology, University of Minnesota Medical School, Minneapolis, MN 55455, USA; Department of Psychiatry, University of Minnesota Medical School, Minneapolis, MN 55455, USA; School of Sport and Exercise Science, Australian Catholic University, 1100 Nudgee Road, Banyo, QLD 4014, Australia

**Author notes:** **Communicating author:** D. N. Lee.

## Abstract

Nervous systems control purposeful movement, both within and outside the body, which is essential for the survival of an animal. The movement control functions of globus pallidus (GP), subthalamic nucleus (STN) and zona incerta (ZI) were analyzed in monkeys reaching for seen targets. Temporal profiles of the hand movements of monkeys and the synchronized flow of electrochemical energy through these basal ganglia were analyzed in terms of a theory of goal-directed movement. Theoretical and empirical analysis indicated: (i) the neural information for controlling movement is the relative-rate-of-change of flow of electrochemical energy in neurons rather than the flow itself; (ii) GP is involved in creating *prospective* electrochemical flow to guide movement; (iii) STN is involved in registering the *perceptual* electrochemical flow monitoring the movement; (iv) ZI is involved in integrating the prospective and perceptual electrochemical flows to power the muscles and thence the movement. Possible implications for PD are discussed.

---

Acting purposefully is vital for the survival of any organism. Controlling purposeful action is the principal function of the nervous system. Purposeful action involves prospective, future-directed guidance of body parts to goals, so they arrive with appropriate momentum. For example, a seabird must move its bill forcefully to spear fish but gently to feed its young; a jaguar must move its feet forcefully when running, but gently when stalking. Purposeful action also requires *perceptually monitoring* the action and *powering the muscles* appropriately. These three essential principles of purposeful action control – prospective guidance, perceptual monitoring and powering muscles – are all enacted by the brain and nervous system.

Here we report an experimental study of the function of basal ganglia in controlling purposeful action. Monkeys moved their hands to a seen goal, while their hand movement and the electrical activity in basal ganglia were synchronously monitored. The study was based on a general theory of action control (Tau/RhoTheory) that was developed from General Tau Theory (Lee et al. 2009), which was inspired by the pioneering theories of Gibson (1966) and Bernstein (1967). Principal tenets of Tau/Rho Theory are as follows:

i. *Action-gaps*. Purposeful movement entails guiding the movement of body parts to goals across *action-gaps.* Action-gaps can be across any physical dimension - e.g., distance when reaching; angle when looking; force when gripping; intra-oral pressure when suckling; pitch, loudness and timbre when vocalizing.
ii. *Tau/Rho* is the primary information used in controlling gaps. *Rho* of a gap equals the *relative rate of closing, or opening* of the gap. *Tau* of a gap equals 1/rho of the gap, which equals the time-to-closing, or time-from-opening of the gap at the current rate of closing, or opening. Thus, *rho* and *tau* of a gap, *X,* equal *Ẋ* / *X* and *X* / *Ẋ* respectively, where the dot indicates the time derivative. Rho and tau are directly perceptible through all known perceptual systems, in contrast with gap size, velocity or acceleration, which are not directly perceptible because they require scaling (Lee 1998).
iii. *Tau/rho-coupling.* This enables the synchronous closing of two gaps, *Y* and *X.* Rho coupling involves keeping rhoY proportional to rhoX, by following the rho-coupling equation:

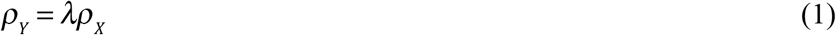 *λ* is the coupling factor, which determines the gentleness (*λ* > 2) or forcefulness (*λ* < 2) of the gap closure. For example, catching a ball gently or forcefully is achievable by keeping the rho of the hand to catching place coupled to the rho of the ball to the catching place, with *λ* > 2 or *λ* < 2. Rho-coupling also features centrally in the following.
iv. *Stimulus-power.* The rho of an action-gap between an external surface element and a sensor is present in the ‘*stimulus-power*’ incident on the sensor, i.e., in the rate of flow of energy from surface element to sensor. In vision, the stimulus-power is electromagnetic, in hearing and echolocating it is mechanical (vibrational), in touch it is mechanical, in smelling it is chemical, in heat-sensing it is thermal, in electrolocating it is electrical. In general, stimulus-power is proportional to the inverse square of the gap between of object and sensor. Thus, the rho of stimulus-power equals half the rho of the action-gap, i.e.,

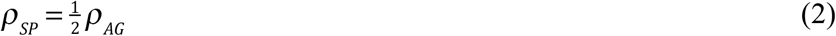
v. *Pain.* When the stimulus-power exceeds the pain threshold, the rho of the stimulus-power, and hence of the action-gap, is registered by nociceptors, which thereby provide information for controlling the closing or opening of a potentially harmful action-gap.
vi. *Neural-power.* At the sensor, or nociceptor, the stimulus-power is converted into *neural-power* (perceptual), which passes into the nervous system. *Neural-power* is the rate of flow of electrochemical energy in the nervous system, along axons and dendrites and across synapses. The general term neural-power is introduced to emphasize that power, rate of flow of energy, is the ubiquitous conveyor of information throughout a nervous system, whatever form neural-power may take and however it may be measured. For example, in neuronal axons in the CNS of mammals neural-power takes the form of a train of action potentials of approximately uniform energy (Kandel *et al.* 2000), which are produced by sodium/potassium pumps injecting bursts of ionic energy at nodes of Ranvier, to speed transmission. Neural-power in axons is often recorded as electrical spike-rate. However, when an axon synapses on a dendrite of another cell the neural-power in the axon can trigger the release of a chemical neurotransmitter, and is thereby converted into neural-power (chemical). Post-synaptically, the neural-power (chemical) becomes neural-power (ionic) in dendrites, which is often recorded as graded synaptic potentials. The neural-power in the dendrites is spatially and temporarily summated in the soma, resulting in an aggregate neural-power in the cell’s axon, which is recorded as a train of action potentials. And so the process continues. Assuming the conversion of stimulus-power into neural-power follows a power law - i.e., neural-power (perceptual) is proportional to stimulus-power raised to some exponent, *α* - it follows that the rho of neural-power (perceptual) equals *α* times the rho of stimulus-power, i.e.,

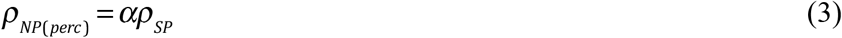
vii. *Prospective neural-power.* To fulfill an organism’s purpose, its nervous system must *prospectively* control the kinematics of closure of action-gaps. In other words, it must set up the pattern of closure in advance. A way of doing this is to rho-couple the action gap to a prospective neural-power gap created in the nervous system. Motion-capture analyses of a range of skilled actions by humans, animals and cells (Lee et al. 2009) have indicated the use of two types of prospective neural-power gap, *G* and *D,* that change, respectively, at constant accelerating or decelerating rates, rather like a ball bouncing on the ground. When the neural-power representation of an action-gap, *AG*, is rho-coupled to a G-type prospective neural-power gap, it follows the rho-coupling equation

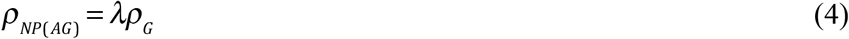

where *λ* is a coupling factor. *ρ_G_* is derived from Newton’s equations of motion, as

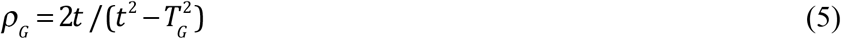

where time *t* runs from 0 to *T_G_*, the duration of the gap-closing movement. An action-gap that follows equations (4) and (5) is said to be *rhoG-guided.* The velocity profile of a rhoG-guided movement is determined by the coupling factor, *λ*. The velocity profile is single-peaked and the temporal position of the peak is determined by the value of *λ* (Fig. 1). When *λ* > 1 the gap-closing movement first accelerates at a varying rate up to a peak velocity and then immediately decelerates at a varying rate to the goal. Gently touching an object with a velocity at contact of zero (as when a seabird feeds its chick) requires *λ* ≥ 2. Hitting something, so that the velocity at contact is positive (as when a seabird spears a fish) requires 0 < *λ* < 2, with lower *λs* producing greater force at impact. *RhoG* and *rhoD* possibly have a gravitational origin. For example, when an animal is running, at every stride it alternates between being in free-fall under gravity and being supported by the ground. As it passes from free-fall to support, mobile masses within cavities of its body, notably the vestibular system in mammals, accelerate at a constant rate downward under gravity relative to the cavities, and so the motion of the mass follows a rhoG function. The opposite occurs when passing from support to free-fall: the masses accelerate at a constant rate upward relative to the cavities, and so follow a rhoD function. Thus, when running, the otolith organs in the vestibular system (Highstein et al., 2004) will be constantly stimulated by rhoG and rhoD functions, which will be transduced into rhoG and rhoD functions of neural-power, which will circulate in the nervous system, which will thus provide material for creating rhoG and rhoD prospective-guides, which could be used in controlling the limbs when running, and in other movements too.

**Fig. 1.**
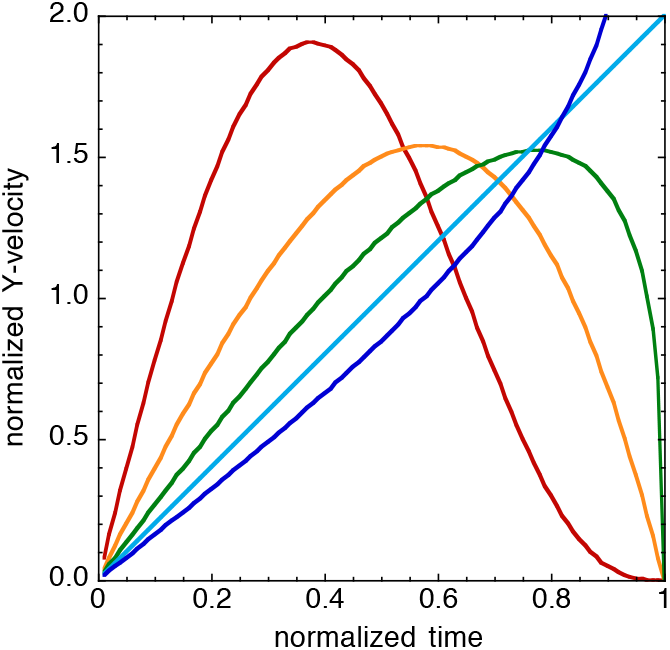
RhoG-guidance of gap *Y* following equation (2): *ρ_Y_* = *λρ_G_*. Effect of the value of coupling factor, *λ*, on the velocity of closure of *Y*: red *λ* = 4; orange, *λ* = 2; green, 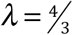 light blue, *λ* = 1; blue, 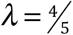. *Y* and movement duration, *T_G_*, are each normalized to 1 for illustration.

There is evidence for rhoG- or rhoD-guidance of movement from experiments spanning a range of skills. (The experiments were reported with *τ_G_* and *τ_D_* as the experimental variables rather than *ρ_G_* and ; however, since, *ρ* = 1/*τ*, the experimental results are directly translatable into *ρ* terms.) The experiments included newborn babies suckling (Craig & Lee 1999); infants catching (van der Meer *et al.* 1994); adults reaching (Lee et al. 1999), controlling gaze (Grealy *et al.* 1999; Lee 2005), intercepting moving objects (Lee *et al.* 2001), putting at golf (Craig *et al.,* 2000), flying aircraft (Padfield 2011), singing and playing music (Schogler *et al.* 2008); hummingbirds aerially docking on a food source (Lee *et al.* 1991; Delafield-Butt *et al.* 2010); unicellular paramecia steering to an electric pole (Delafield-Butt *et al.* 2012). There is also evidence for rho (tau) at a neural level - in the brains of locusts (Rind & Simmons 1999), pigeons (Sun and Frost 1998), monkeys (Merchant et al. 2004; Merchant & Georgopoulos, 2006) and humans (Field & Wann, 2005; Tan et al., 2009; van der Weel *et al.* 2009).

We investigated neural processes underpinning rhoG-guidance by analyzing single unit recordings from external and internal globus pallidus (GPe, GPi), subthalamic nucleus (STN) and zona inserta (ZI), when monkeys were moving their hand to a goal along a straight track. These basal ganglia are thought to be involved in sensorimotor control (Mitrofanis 2005; Fasano et al. 2015; Takamitsu & Yamomoto 2015)). There is also evidence of temporal coherence in basal ganglia during voluntary movement (Talakoub et al. 2016). However, how the temporal pattern of neural-power in basal ganglia relates to the temporal pattern of voluntary movement has not been studied. This was our aim.

## RESULTS

### Relation between neural-power and hand movement

On each trial, the neural-power in GPi, GPe, STN and ZI was measured as spike-rate, the rate of flow of action potentials. The neural-power was time locked onto the start of the hand movement and time-normalized into bins of duration 0.05 times the duration of the hand movement on that trial; i.e., into bins of duration 0.05 movement-time units (mtus). The mean and standard error of the time-normalized neural-power time series were then computed across all the trials in each ganglia under study. The black lines in Figure 3A show the means and standard errors of the time-normalized neural-powers. The coloured lines show these neural-powers smoothed with a Gaussian filter, sigma 0.1 mtu.

Even though the means and standard errors of the time-normalized neural-powers (black lines in Fig. 2A) were computed across many different neurons (214 in GPe, 86 in GPi, 33 in STN, 68 in ZI) and in three monkeys (Methods), the standard errors (se) were remarkably small. As a fraction of the maximum value of neural-power, the mean ± se of the standard errors of mean neural-power were: 0.00157 ± 0.00001 (GPe), 0.00108 ± 0.00001 (GPi), 0.00123 ± 0.00003 (STN), 0.00121 ± 0.00001 (ZI). These small standard errors (possibly related to the monkeys’ hand movements on the task being well practiced) strongly suggest that the average duration-independent neural-power (black lines in Fig. 2A) measures a basic time-invariant of the hand movement, and thereby provides a measure of the average neural activity (across neurons in a tract) underpinning the hand movement.

**Fig. 2.**
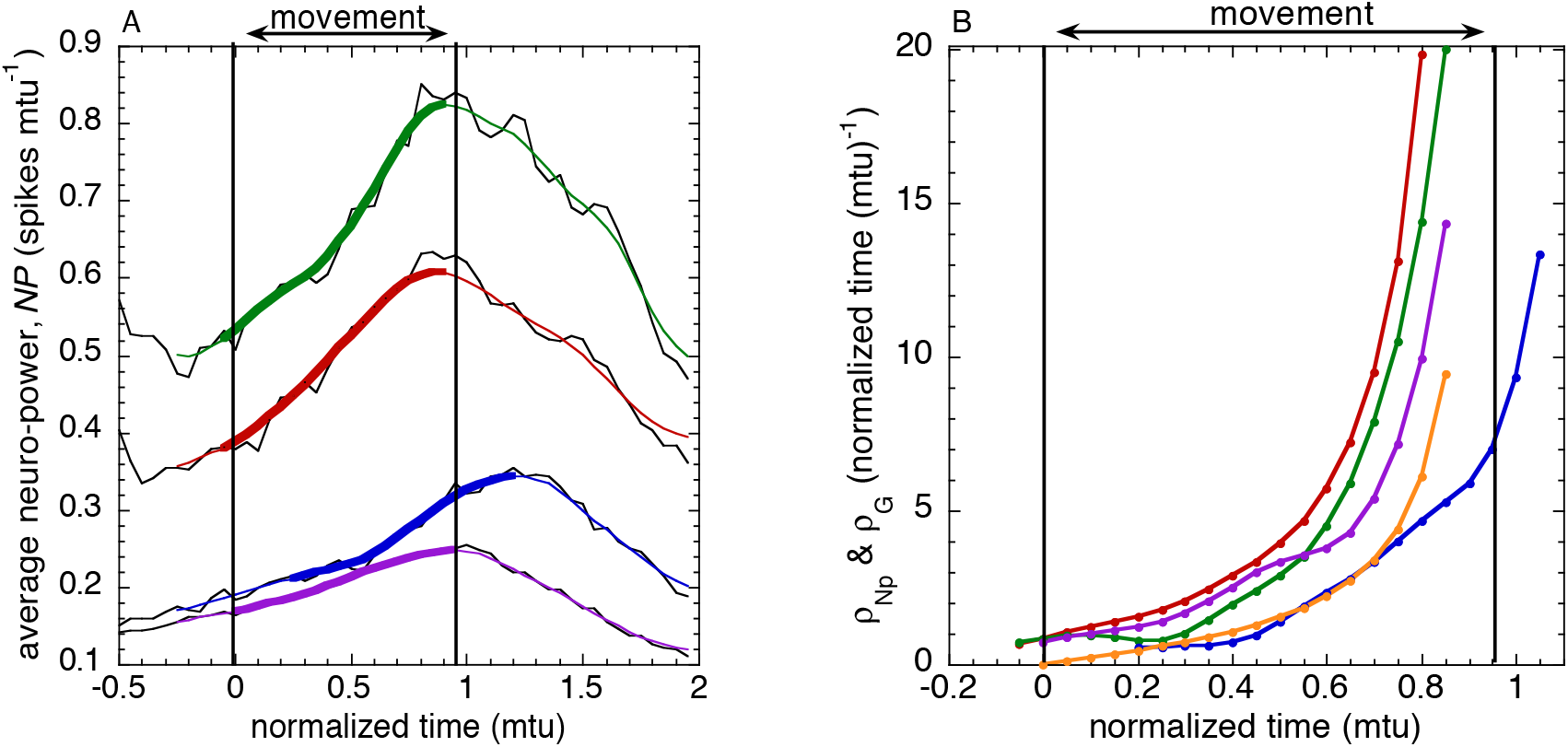
(A) Neural-power profiles during hand movement. The neural-power profiles were measured as spike-rate and were time-normalized with respect to hand movement time and time-locked to it. Black lines: means of unsmoothed neural-power. Coloured lines: Gaussian (sigma 0.1 mtu) smoothed values of GPe (red), GPi (green), STN (blue), ZI (purple). Thicker coloured lines: the sections of the neural-power profiles that were of the same duration as the hand movement (1 mtu), and the *ρ* s of which were the most highly correlated with *ρ_Mov_*, the *ρ* of the hand movement. (B) The *ρ* s of the neural-power sections in (A), together with *ρ_G_* (orange line).

The sections of the mean time-normalized neural-power profiles (Fig.2A) that were most strongly rho-coupled to the hand movement were determined by computing, for each time-normalized neural-power profile, the *rho*s of the neural-power sections of duration 1 mtu (movement time unit) that ended at each point in the neural-power profile. The rho of each of these sections of neural-power, *ρ_Np_*, was then linearly regressed on *ρ_Mov_*, the rho of the movement up to the goal position, and the section of the neural-power profile that yielded the highest *r*^2^ was considered to correspond to the hand movement. These sections of the neural-power profile were found to end at the peak mean time-normalized neural-power in each ganglia studied. In GPe and GPi, the peak occurred 0.05mtu (on average about 25ms) before the movement ended. In ZI it occurred just as the movement ended. In STN it occurred 0.2mtu (on average about 100ms) *after* the movement ended. Fig. 2B shows *ρ_NP_* (the rho of the section of the neural-power profile up to the peak neural-power) for the thick-lined sections of the neural-power profiles corresponding to the hand movement (Fig. 2A). Also shown is rhoG (orange line), the rho of the predicted prospective neural-power.

### Coupling *ρ*_*NP*(*GPe*)_ onto *ρ_G_*

The degree to which the *ρ* of neural-power (*ρ_NP_*) was rho-coupled onto *ρ_G_* was measured by plotting *ρ_NP_* against *ρ_G_* for each of the four ganglia, and calculating the linear regressions forced to pass through the origin (since *proportionate* coupling was predicted). The regression lines are shown in Fig. 3A. The regression coefficients are given in Table 1. The *R^2^* values, the coefficients of determination, measure the strengths of the rho-couplings, i.e., the proportion of variance in *ρ_NP_* accounted for by the rho-couplings. The strength of the rho-coupling of *ρ*_*NP*(*GPe*)_ onto *ρ_G_* was very high (*R*^2^ = 0.996, compared with a maximum possible value of 1.000). This strongly suggests that GPe was involved in creating the prospective-guiding function, *λρ_G_*. The slopes of the regressions (Table 1) measure the *λ* coupling factors. For *ρ*_*NP*(*GPe*)_ on *ρ_G_*, *λ* was 2.13, which indicates that the neural-power in GPe approached its peak value gently, stopping when it got there (c.f. Fig.1).

**Table 1.** Degrees of rho-coupling (R^2^ of linear regressions through origin). In parentheses, rho-coupling factors (the regression slopes). Red, R^2^ ≥0.990; purple, R^2^ ≥0.980. Rows, dependent variables; columns, independent variables (e.g., *ρ*_*NP*(*GPe*)_ = 2.13*ρ_c_*)

|  |  |  |  |  |  |  |
| --- | --- | --- | --- | --- | --- | --- |
| $\rho_{NP(GPe)}$ | 0.996<br>(2.13) | | 0.990<br>(1.312) | 0.988<br>(1.316) | 0.969<br>(1.343) | 0.965<br>(0.278) |
| $\rho_{NP(GPi)}$ | 0.986<br>(1.615) | 0.989<br>(0.758) | | 0.974<br>(0.996) | 0.963<br>(1.019) | 0.954<br>(0.219) |
| $\rho_{NP(ZI)}$ | 0.973<br>(1.604) | 0.987<br>(0.755) | 0.970<br>(0.988) | | 0.980<br>(1.020) | 0.982<br>(0.214) |
| $\rho_{NP(STN)}$ | 0.960<br>(1.559) | 0.965<br>(0.732) | 0.956<br>(0.961) | 0.980<br>(0.970) | | 0.989<br>(0.212) |
| $\rho_{Mov}$ | 0.984<br>(7.358) | 0.969<br>(3.546) | 0.948<br>(4.453) | 0.975<br>(4.609) | 0.986<br>(4.687) | |
| | $\rho_G$ | $\rho_{NP(GPe)}$ | $\rho_{NP(GPi)}$ | $\rho_{NP(ZI)}$ | $\rho_{NP(STN)}$ | $\rho_{Mov}$ |

**Fig. 3.**
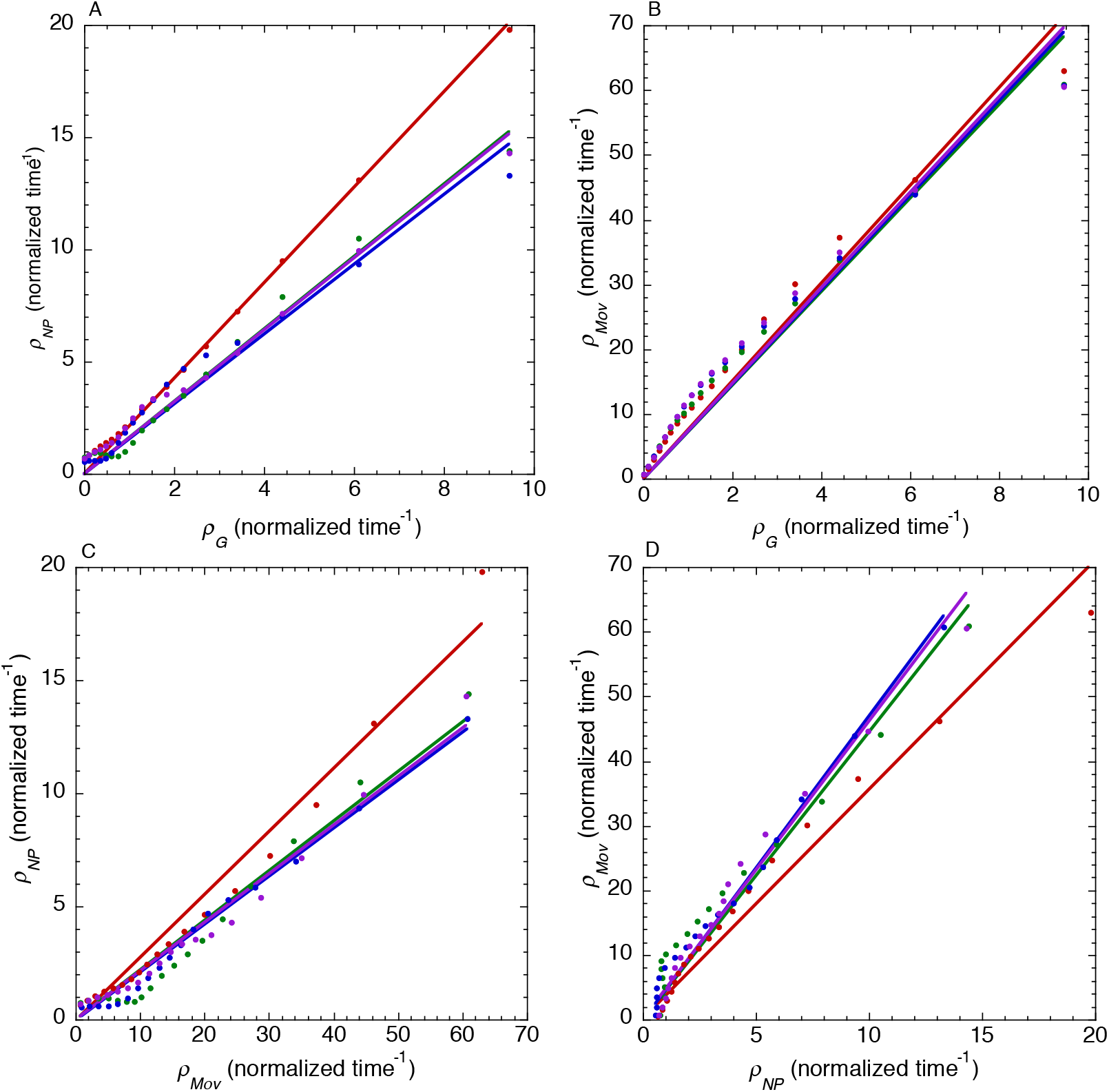
Rho-coupling regressions through origin. (A) Rho of neural-power (*ρ_NP_*) on hypothesized prospective rho (*ρ_G_*). (B) Rho of hand movement, *ρ_Mov_*, on *ρ_G_*. (C) *ρ_NP_* on *ρ_Mov_* · (D) *ρ*_*Mov*_ on *ρ_NP_*.

### Coupling *ρ_Mov_* onto *ρ_G_*

The degree to which the *ρ* of the movement (*ρ_Mov_*) was rho-coupled onto *ρ_G_* was measured by plotting the average *ρ_Mov_* across the four ganglia studied and calculating the linear regressions through the origin (since *proportionate* coupling was predicted). The regressions are shown in Fig. 3B. The regression coefficients are given in Table 1. The *R*^2^ for the coupling of *ρ_Mov_* onto *ρ_G_* was 0.984. This indicates that *ρ_Mov_* followed *ρ_G_* quite closely. The coupling factor was 7.358, indicating that the hand approached the target very gently, slowing down quite early to stop at it (c.f. Fig. 1).

### Coupling *ρ_NP_* onto *ρ_Mov_*

The degree of coupling of *ρ_NP_* onto *ρ_Mov_* was measured by plotting *ρ_NP_* against *ρ_Mov_* for each ganglion and calculating the linear regressions through the origin (since proportionate coupling was predicted). The regressions are shown in Fig. 3C. The regression coefficients are given in Table 1. The neural-power section in STN that corresponded to the hand movement started 0.2 mtu *after* the hand movement started. The highest *R^2^* (0.989) was for STN. Taken together these findings suggest that STN was involved in the perceptual monitoring of *ρ_Mov_*, after a perceptual delay of 0.2 mtu (about 100 ms on average).

### Coupling *ρ_Mov_* onto *ρ_NP_*

The degree of coupling of *ρ_Mov_* onto *ρ_NP_* was measured first by plotting *ρ_Mov_* against *ρ_Np_* for each ganglion studied and calculating the linear regressions through the origin. The regressions are shown in Fig. 3D. The regression coefficients are given in Table 1. The two highest R^2^ were for *ρ*_*NP*(*STN*)_ (0.986) and *ρ_G_* (0.984). This suggested to us that *ρ*_*NP*(*STN*)_ and *ρ_G_* might have been jointly influencing *ρ_Mov_*. In particular, that ZI might receive movement-adjusting *ρ* information from STN and GPe (since *ρ*_*NP*(*GPe*)_ was very strongly coupled to *ρ_G_*; *R^2^* = 0.996) and transmit this information to the muscles, and thence to the hand movement. To investigate this possibility we defined *ρ*_*NP*(*ZI*)_(*t*) = *f*(*ρ*_*NP*(*GPE*)_(*t*),*ρ*_*NP*(*STN*)_(*t*)), and then sought to determine the function *f* from the experimental data. Taking into account the observed perceptual delay of 0.2 mtu, when STN could not have registered the hand movement, we hypothesized that, for t = 0 to 0.2 mtu,

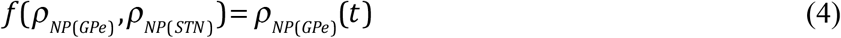

and for t = 0.2 to 1.0 mtu,

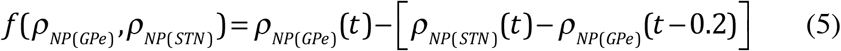

where [*p*_*NP*(*STN*)_(*t*)–*ρ*_*NP*(*GPe*)_(*t* – 0.2)] is the deviation of the movement as perceived from its prospective rho course. To test the hypothesis we computed the linear regression (through the origin) of *ρ*_*Np*(*ZI*)_ on *f*(*ρ*_*NP*(*GPe*)_’*ρ*_*NP*(*STN*)_) for t = 0 to 1.0 mtu. The regression yielded an *R^2^* of 0.993 and a coupling factor of 0.740. Thus, the hypothesis was strongly supported.

**Fig. 4.**
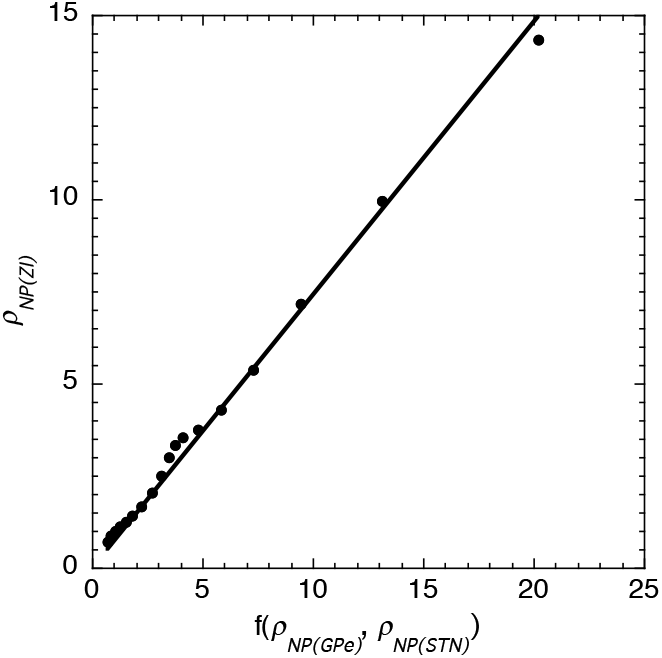
*ρ* of neural-power in ZI. *ρ*_*NP*(*ZI*)_, is plotted against the *ρ* input to ZI, *f*(*ρ*_*NP*(*GPe*)_,*ρ*_*NP*(*STN*)_); see equations (4) and (5). The linear regression line was forced to pass through the origin to test predicted *proportional ρ* - coupling. The regression equation was 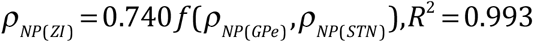.

## Discussion

We have argued that animals prospectively control their actions by using, as information in the nervous system, the relative-rate-of-change, *ρ*, of neural-power (i.e., the rate of flow of electrochemical energy through neurons). A principal argument that *ρ* is the fundamental informational variable for controlling action is that the *ρ* of the distance gap that needs to be closed to achieve a controlled action is directly perceptible, whereas the distance itself, or any of its time derivatives, are not directly perceptible.

The results of our analysis of the electrical activity in basal ganglia GPe, GPi, STN and ZI of monkeys reaching to seen targets is consistent with the idea that GPe is implicated in *prospectively guiding* the *ρ* of a movement using the prospective neural-power function, *ρ_G_*; that STN is implicated in the *perceptual monitoring* of the movement; and that ZI is implicated in integrating (Sherrington 1961, Branco et al 2010, 2012) the *ρ* of *prospective* neural-power from GPe and the *ρ* of *perceptual* neural-power from STN to create the *ρ* of *enacting* neural-power at ZI. The *ρ* of the *enacting* neural-power both informationally and physically (after amplification, e.g., with ATP) powers the *ρ* of *muscular-power* (the rate of flow of energy into the muscles) and this powers the *ρ* of the *action-power* of the movement. Thus the *ρ* of power has come full circle.

Perhaps it is not too surprising that the basal ganglia should be implicated in the three fundamental neural functions controlling action - prospecting, perceiving and powering. After all, basal ganglia are phylogenically ancient in vertebrates - including those, like lamprey, who lack cerebral cortices (Grillner 2003) – and all vertebrates control their movements purposefully in order to survive.

### Parkinson’s Disease

The motor symptoms of Parkinson’s Disease - tremor, rigidity, bradykinesia, freezing, and dysarthria - are generally considered to involve malfunction in the basal ganglia (Moustafa, et al., 2016). However, the electrophysiological functions in the basal ganglia that are affected are unknown (Ellens & Leventhal, 2013). The present study could cast some light on the issue. Many movements are affected in Parkinson’s disease, but some are relatively unaffected - the so-called paradoxical movements. For example, catching a moving ball can be easier than reaching for a stationary one, and walking downstairs can be easier than walking across a featureless floor. The difference in ease of performance could be related to the type of information being used. The information for catching a moving ball or walking downstairs is largely perceptual, from the optic flow field at the eye; whereas, when reaching for a stationary ball or walking across a featureless floor, action-control is more reliant on prospectively-guiding information created in the nervous system. Since our results indicate that GPe is strongly implicated in generating prospectively-guiding *ρ_G_* information, it is possible that movement disorders in Parkinson’s – tremor, rigidity, bradykinesia, freezing, and dysarthria - which all involve poor prospective coordination of muscles – may be due, in part at least, to GPe dysfunction.

### Where next?

A principal function of any nervous system is controlling bodily actions. If, as our results suggest, the common informational currency in basal ganglia for controlling actions is the *ρ* of neural-power, then it is likely that the same informational currency is used throughout the nervous system when actions are being controlled (otherwise a Tower of Babel situation would prevail). This idea could be tested in humans, for example, by using high temporal resolution MEG (Tan et al., 2009). If verified, rho theory might then be used to analyze normal function in the nervous system, and also reveal regions of the brain where there is dysfunction in the transmission of information for guiding movement.

Rho theory might also be useful in investigating how other organisms control their movements. For example, Delafield-Butt *et al.* (2011) have obtained motion-capture evidence that unicellular paramecia guide their movement to an electric pole using the *ρ_D_* function as a guide; Strausfeld and Hirth (2015) have suggested that the central complex in an insect’s brain is homologous with the basal ganglia in animals, and so might have similar control functions; and Darwin and Darwin (1880) and Masi *et al.* 2009 have suggested that the transition zone in the roots of plants is implicated in neural control of movement. It would be of value to investigate to what extent the *ρ* of neural-power is used generally by organisms in controlling their actions.

### Methods

We analyzed neural and movement data from three rhesus monkeys in four experiments. In each experiment there were two horizontal rows of 128 LEDS, 32 cm long, one above the other. The monkeys were trained to move a handle to the left or right to line up an LED on the lower row with a target LED on the upper row. The handle was constrained to run along a track 32 cm long. The position of the handle was recorded every 10 ms. The movements averaged 525 ms and 248 mm. Neural electrical activity was recorded extra-cellularly with microelectrodes on separate occasions from five hemispheres of three monkeys: from 214 cells in the arm area of globus palidus external (GPe), 86 cells from globus pallidus internal (GPi), 33 cells in the subthalamic nucleus (STN) and from 75 cells in zona incerta (ZI). Details of the procedures for the experiments are given in DeLong et al. (1985) and Crutcher et al. (1980)

### Data analysis

The data for GPe, GPi, STN and ZI were first assembled into *unit records,* which comprised the hand position data and neural data recorded on a single reach-to-target trial. Using the index *i* to refer to a unit record, and the index *j* to refer to a 10 ms time sample, a unit record comprised (i) a hand position time series, *x_ij_*, where *x* is the coordinate of the handle, and (ii) a neural spike-density time series, *n_ij_*, where *n_i,j_* is the number of neural spikes in the *jth* 10 ms time bin of the *ith* unit record. Then, for each unit record (i) to minimize noise in the data, the *x_i,j_* time series of the hand were smoothed, with a Gaussian filter with time constant sigma of 30 ms, and a cutoff frequency of 6 Hz, yielding the smoothed time series *X_i,j_*; (ii) the *X_i,j_* time series were numerically differentiated with respect to time to yield the time series 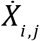; (iii) the movement time, *MT_i_*, for the *ith* unit record was calculated as the difference between the ‘hand-start’ and ‘hand-end’ times, defined respectively as the first sample time after the speed of the hand rose above and the last sample time before it fell below 5% of the peak speed on that recording. *MT_i_* averaged 525ms; (iv) each hand/target action-gap time series, *M_i,j_*, was computed as the distance between the position of the handle at each sample time, *j*, and its position at the end of the movement.

A unit record was accepted for further analysis providing the neuron became ‘active’ between the stimulus light going on and the hand starting to move; i.e., providing there were five or more consecutive 10 ms bins in the time series which were of value greater than three standard deviations above the mean value of *n_i,j_* during the 500 ms preceding the stimulus light. This criterion was applied to exclude normalized unit records where there was no evidence of the neural spike-rate being related to the hand movement. In the records satisfying the criterion (i) the members of the hand/target gap time series, *M_i,j_*, were numerically differentiated with respect to time to yield the time series 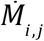; (ii) the time series, *ρ_i,j_*, of the hand/target gap, was calculated using the formula 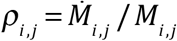; (iii) the time series 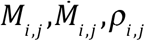 were time-normalized to yield the time series 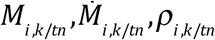. Time-normalization entailed apportioning the data in each time series into equal time bins of width (*MT_i_*/20) s, or 0.05 mtu (movement time units), so that for the movement of the hand between the start time and the end time the index, *k*, ran from 0 to 19, and normalized time ran from 0 mtu to 0.95 mtu. For the *n_i,k/tn_* time series the index *k* ran from −40 through 0 (when the hand movement started) to +39, and the corresponding normalized time ran from −2.00 mtu through 0 (when the hand movement started) to +1.95 mtu. Normalizing time in this way meant that the normalized duration of the hand movement remained the same across the normalized records, enabling the average time-normalized spike-density function, which was assumed to be coupled in *relative* time to the hand movement, to be measured as the mean time series, *n*_•,*k*/*t*/*av*_ of the *n*_*i*,*k*/*tn*_ time series, for *k* = −40, −39,….39, and normalized time running from −2.00 mtu to 1.95 mtu. The values of *n*_•,*k*/*t*/*av*_ were computed separately for the GPe, GPi, STN and ZI time-normalized records. The time series *n*_•,*k*/*t*/*av*_ were also averaged separately across the records, yielding the mean time series, 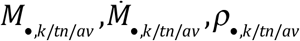 for *k* = 0, 2,….19, and normalized time running from 0 mtu to 0.95 mtu.

To enable statistical comparisons to be made, the same analysis was performed on 1000 samples, drawn with replacement from each of the normalized records. This resulted, for each brain area, in 1000 sets of average time series 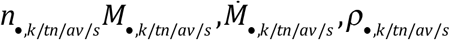, for s = 1, 2,…1000.

### Summary: time series used in the analysis

In summary, the time series used in analyzing the results were 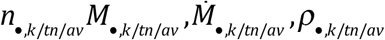. In the main text and figures these average normalized-time time series are, for conciseness, designated respectively as *n* (the average time-normalized spike density function, which is taken as the measure of average *neural-power*), *M* (the average time-normalized action-gap), *Ṁ* (the time derivative of the average time-normalized action-gap), and *ρ_M_* (the *ρ* of the average time-normalized action-gap). These time series are graphed in the figures for normalized time extending from −2 mtu to +2 mtu in steps of 0.05 mtu, which corresponds to *k* = 1, 2,………. 80.

## Acknowledgements

The research was supported by grants from American Legion Brain Sciences Chair, BBSRC, EU NEST-ADVENTURE, Leverhulme Trust, US Public Health Service Grant PSMH48185, US Department of Veterans Affairs.

## References

Bernstein NA (1967) The co-ordination and regulation of movements. Oxford: Pergamon

Branco, T., Clark, B.A., Hausser, M. (2010) Dendritic discrimination of temporal input sequences in cortical neurons. Science, 329, 1671–1675

Branco, T. & Hausser, M. (2011). Synaptic integration gradients in single cortical pyramidal cell dendrites. Neuron, 69, 885–892.

Craig CM, Delay D, Grealy MA, Lee DN (2000) Guiding the swing in golf putting. Nature 405: 295–296

Craig CM, Lee DN (1999) Neonatal control of nutritive sucking pressure: evidence for an intrinsic tau-guide. Exp Br Res 124: 371–382

Crutcher, M.D., Branch, M.H., DeLong, M.R. and Georgopoulos A.P. (1980) Activity of zona incerta neurons in the behaving primate. Soc. Neurosci. Abstr. 6: 676.

Darwin C, Darwin F (1880) The power of movement in plants. London: Murray

Delafield-Butt, J. T., Galler, A., Schögler, B. & Lee, D. N. (2010). A perception-action strategy for hummingbirds. Perception, 39, 1172–1174.

Delafield-Butt, J. T., Pepping, G-J., McCaig, C. D., Lee, D. N. (2012). Prospective guidance in a free-swimming cell. Biological Cybernetics, 106, 283–293.

DeLong, M. R., Crutcher, M. D. & Georgopoulos, A. P. (1985). Primate Globus Pallidus and Subthalamic Nucleus: Functional Organization, J. Neurophysiol. 53, 530–543.

Ellens, D.J., Leventhal, D.K. (2013). Electrophysiology of basal ganglia and cortex in models of Parkinson disease. J. Parkinsons Dis., 3(3), 241–254.

Fasano, A., Aquino, C.C., Kraus, J.K., Honey, C.R. Bloem, B.R. (2015). Axial disability and deep brain stimulation in patients with Parkinson disease. Nature Reviews Neurology, 11, 98–110.

Field, D.T., Wann, J.P. (2005). Perceiving time to collision activates the sensorimotor cortex. Current Biology, 15, 453–458.

Georgopoulos, A.P., DeLong, M.R. & Crutcher, M.D. Relations between parameters of step-tracking movements and single cell discharge in the globus pallidus and subthalamic nucleus of the behaving monkey. J. Neurosci. 3, 1586–1598 (1983).

Gibson JJ (1966) The senses considered as perceptual systems Boston: Houghton Mifflin.

*Grealy, M. A., Craig, C. M. & Lee, D. N. (1999). Evidence for on-line visual guidance during saccadic gaze shifts. Proceedings of the Royal Society of London B, 266, 1799–1804.

Grillner, S. (2003). “The motor infrastructure: From ion channels to neuronal networks”. Nature Reviews Neuroscience. 4 (7): 573–586.

Highstein, S. M., Fay, R. R. Popper, A. N. eds. (2004). The vestibular system. Berlin: Springer.

Kandel, E. R., Schwartz, J. H. & Jessell, T. M. Eds. (2000). Principles of Neural Science 4th Edition. New York: McGraw-Hill.

Lee DN (1998) Guiding movement by coupling taus. Ecol Psychol 10: 221–250.

Lee DN (2005) Tau in action in development. In Rieser JJ, Lockman JJ, Nelson CA (eds) Action as an organizer of learning and development. pp 3–49 Hillsdale NJ: Erlbaum.

Lee, D. N., Bootsma, R. J., Frost, B. J., Land, M. & Regan, D. (2009). General Tau Theory: evolution to date. Special Issue: Landmarks in Perception. Perception, 38, 837–858.

Lee DN, Craig CM, Grealy MA (1999) Sensory and intrinsic coordination of movement. Proc R Soc Lond B 266: 2029–2035.

Lee, D. N., Georgopoulos, A. P., Clark, M. J. O., Craig, C. M., & Port, N. L. (2001). Guiding contact by coupling the taus of gaps. Experimental Brain Research, 139, 151–159.

Lee, D. N., Reddish, P. E. & Rand, D. T. (1991). Aerial docking by hummingbirds. Naturwissenschaften, 78:, 526–527.

Masi E, Ciszak M, Stefano G, Renna L, Azzarello E, Pandolfi C, Mugnai S, Baluska F, Arecchi FT, Mancuso S (2009) Spatiotemporal dynamics of the electrical network activity in the root apex. PNAS 106: 4048–4053

Merchant H, Battaglia-Mayer A, Georgopoulos AP (2004) Neural responses during interception of real and apparent circularly moving stimuli in motor cortex and area 7a. Cerebral Cortex 14: 314–331

Merchant H, Georgopoulos AP (2006) Neurophysiology of perceptual and motor aspects of interception. J Neurophysiol 95: 1–13

Mitrofanis, J. (2005). Some certainty for the “zone of uncertainty”? Exploring the function of the zona incerta. Neuroscience 130 (1): 1–15.

Moustafa, A.A., & Chakravarthy, S., Phillips, J.R., Gupta, A., Keri, S., Polner, B., Frank, M.J., Jahanshahi, M. (2016). Motor symptoms in Parkinson’s disease: A unified framework. Neuroscience and Biobehavioral Reviews 68: 727–740.

Padfield, G.D. (2011). The tau of flight control. The Aeronautical Journal, 115 No 1171, 521–556

Rind FC, Simmons PJ (1999) Seeing what is coming: building collision-sensitive neurons. TINS 22: 215–220

Schogler B, Pepping G-J, Lee D. N. (2008) TauG-guidance of transients in expressive musical performance. Exp Br Res 198: 361–372.

Sherrington CS (1961) The integrative action of the nervous system. New Haven: Yale UP

Strausfeld, N. J. & Hirth, F. (2013). Deep homology of arthropod central complex and vertebrate basal ganglia. Science, 340, 157–161.

Sun H, Frost JF (1998) Computation of different optical variables of looming objects in pigeon nucleus rotundus neurons. Nature Neurosci 1: 296–303

Takamitsu C. & Yamomoto, F. (2015). Deep brain stimulation for Parkinson’s disease: recent trends and future direction. Neurologia medico-chirurgica, 55(5), 422–431.

Talakoub, O., Neagu, B., Udupa, K., Tsang, E. Chen, R., Popovic, M.R. & Wong, W. (2016). Time course of coherence in the human basal ganglia during voluntary movements. Nature Scientific Reports, | 6:34930 | DOI: 10.1038/srep34930

Tan, H-R. M., Leuthold, A. C., Lee, D. N., Lynch, J. K. & Georgopoulos, A. P. (2009) Neural mechanisms of movement speed and *tau* as revealed by magnetoencephalography. Experimental Brain Research, 195, 541–552.

Van der Meer, A.L.H., Van der Weel, F.R. & Lee, D.N. (1994). Prospective control in catching by infants. Perception, 23, 287–302.

Van der Weel, F.R. & Van der Meer, A.L.H (2009). Seeing it coming: Infants’ brain responses to looming danger. Naturwissenschaften. 96: 1385–1391

